# Sex decreases the pleiotropic costs of local adaptation

**DOI:** 10.1101/2025.03.28.646016

**Authors:** Shreyas V. Pai, Parris T. Humphrey, Camille Simonet, Katya Kosheleva, Anurag Limdi, Michael M. Desai

## Abstract

Understanding the evolutionary mechanisms that maintain sex throughout nature despite its substantial direct costs is a longstanding challenge in biology. Previous work has shown that sexual recombination provides a key advantage in speeding adaptation, in part by separating beneficial mutations from deleterious hitchhikers. However, these earlier studies have focused on the effects of sex in a constant environment. Here, we show that recombination also provides a key advantage in fluctuating conditions, promoting the evolution of generalist phenotypes by reducing the pleiotropic costs of local adaptation. Using laboratory evolution in S. cerevisiae as a model system, we show that hitchhiking genetic load leads to pleiotropic costs and hence specialization in response to local adaptation in asexual but not in sexual lineages. This provides the first direct evidence that sex can be maintained over longer evolutionary timescales because it enables lineages to persist in the face of environmental change.

## Main Text

Explaining why sex is pervasive across eukaryotes, despite its substantial direct costs, is a longstanding question in evolutionary biology. Extensive prior theoretical and experimental work has highlighted mechanisms by which sex can speed adaptive evolution, for example by bringing together beneficial mutations that occur in different genetic backgrounds, or by purging hitchhiking deleterious load (*1*–*6*). However, this work has focused on how sex speeds adaptation in a single environment [but see (*7*)]. Here, we analyze a novel potential benefit of sex when environmental conditions fluctuate across time or space: recombination can promote generalism by alleviating the pleiotropic costs of local adaptation. Specifically, sex can decouple locally adaptive mutations from other hitchhiking variation which may have negative effects in other environments, providing an evolutionary advantage over longer timescales.

This potential advantage of sex is related to the “Red Queen” hypothesis, which posits that recombination speeds adaptation in the face of changing selection pressures inherent in arms-race dynamics, for example in host-pathogen coevolution (*8*–*10*). However, while the red queen focuses on how sex speeds adaptation to a rapidly changing target, here we analyze the role of sex in mitigating negative tradeoffs across fluctuating conditions. We expect the strength of this effect to depend on the genetic architecture of pleiotropy. If neutral or mildly deleterious hitchhiking variation tends to be costly in other conditions, purging this variation via recombination will mitigate the tendency of local adaptation to lead to specialization. On the other hand, if specialization results from direct effects of locally adaptive alleles [i.e. antagonistic pleiotropy (*11*)], then sex will not have this effect.

To test for this potential effect of sex in promoting generalism, we describe here the first comparison of the pleiotropic costs of adaptation in sexual versus asexual populations, using laboratory evolution in the budding yeast *S. cerevisiae* as a model system. We evolve both sexual and asexual populations in rich laboratory media, and measure the effect of regular bouts of recombination on the rate and pleiotropic consequences of adaptation. By comparing both molecular and phenotypic evolution in sexual versus asexual populations, we analyze how recombination mitigates the costs of local adaptation and promotes the evolution of generalism.

## Results

We established 17 replicate sexual and 18 replicate asexual *S. cerevisiae* populations from a single laboratory strain, and propagated them independently for 960 generations of laboratory evolution in rich media (YPD) using a standard batch culture protocol (Methods). Sexual populations were propagated by interspersing asexual mitotic growth with regular bouts of recombination every 120 generations, using a system we established in earlier work which ensures that sexual and asexual populations are otherwise maintained as identically as possible (Fig. S1, Methods) (*5, 12*).

### Sex increases the rate and repeatability of adaptation, and decreases its pleiotropic costs

After 960 generations of adaptation, we measured the competitive fitness of each population in the ‘home’ environment, YPD. We find that sexual populations increased in fitness by 8% in this home environment, compared to 5.7% in asexuals. Thus, consistent with previous work, sex increases the speed of adaptation (Fig. 1a; Generalized Wilcoxon test, p < 0.001).

**Fig. 1.**
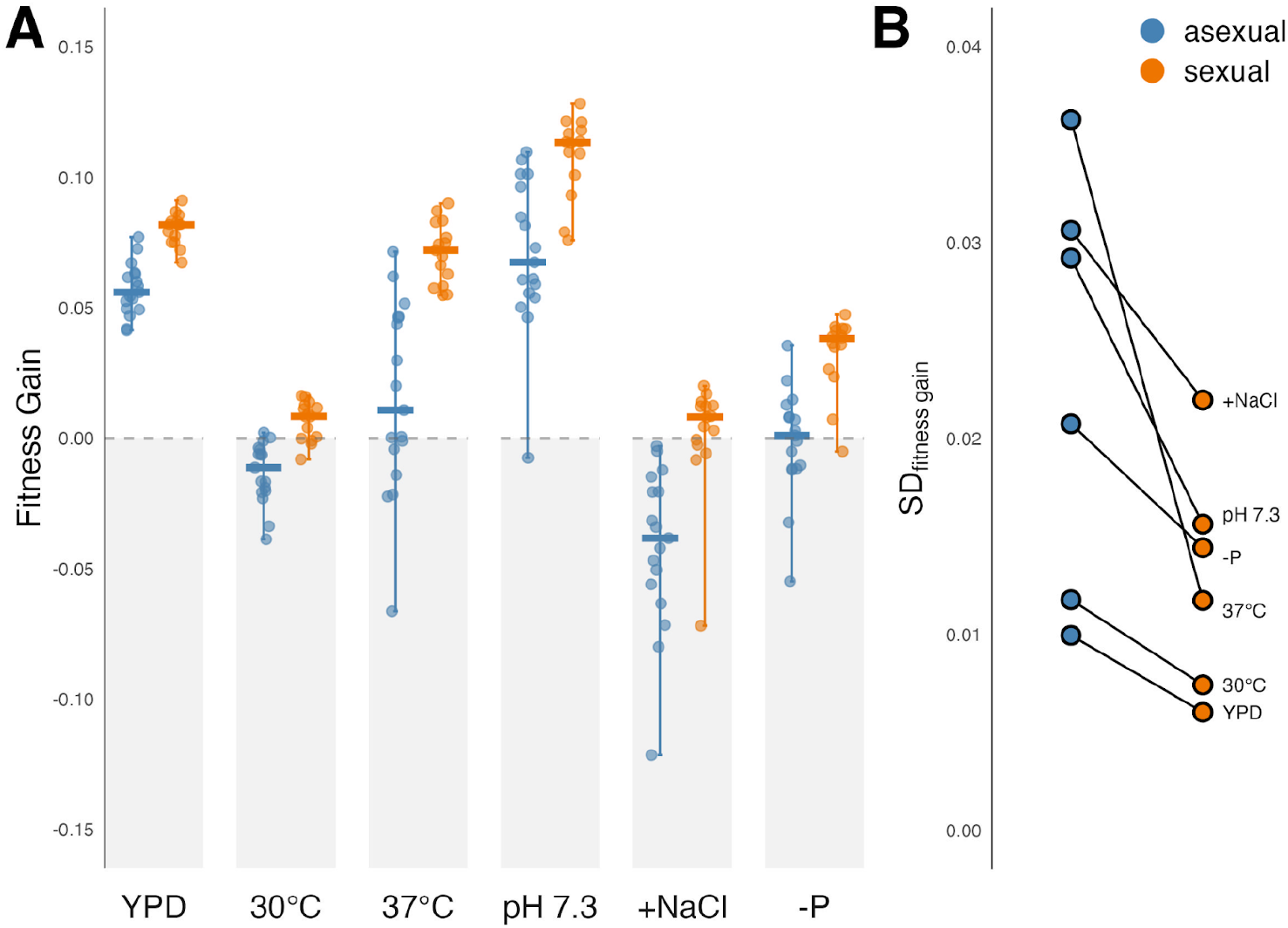
Recombination increases fitness gain, repeatability, and decreases pleiotropic costs of adaptation. **(A)** Distribution of fitness changes relative to the ancestor across home (YPD) and away environments (SC 30°C, 37°C, pH 7.3, +NaCl, -phosphate), with each independent sexual or asexual population represented. Horizontal bars indicate median and vertical lines indicate range. **(B)** Standard deviation in fitness across replicate populations in asexual compared to sexual populations in home and each away environment.

We next measured pleiotropic effects of this adaptation on the fitness of each evolved population in five ‘away’ environments (SC with perturbation: 30°C, 37°C, +0.2 M NaCl, buffered pH 7.3, phosphate limitation). We find that sex also mitigates the costs (and in some cases enhances the benefits) of local adaptation to the home environment: sexual lines exhibit an average fitness advantage of between 2-5.6% over asexuals in each of the five away conditions (Fig. 1a; Generalized Wilcoxon test with Benjamini-Hochberg correction, all p < 0.001). In asexual populations, adaptation to YPD often results in fitness declines in the away environments (Fig. 1a), indicating specialization relative to the ancestor. In contrast, the pleiotropic fitness effects of adaptation are very rarely deleterious for sexual lines (Fig. 1a), even though sexual lines adapted more rapidly to the home environment. Consistent with this, we also find that in both sexuals and asexuals, populations that are more fit at home are also more fit away (Fig. S2, permutation test of pairwise Spearman correlations, mean ρ = 0.476 for asexuals and 0.386 for sexuals, both p < 0.001), suggesting that antagonistic pleiotropy is not the major cause of specialization.

A corollary theoretical prediction is that sex should narrow the range of evolutionary outcomes seen among independent lines evolving in the same environment, because recombination alleviates stochastic effects of clonal interference among beneficial lineages and reduces the effect of genetic draft that would otherwise carry neutral and mildly deleterious mutations to high frequencies. However, prior experimental work has found mixed support for this effect (*5, 7, 12, 13*). Here, consistent with the theoretical expectation, we find that the range of fitness outcomes was consistently more variable among asexual populations than among sexual populations across both home and away environments (Fig. 1b; paired t-test of σ values, t = 3.19, p = 0.02).

### Sex narrows the outcomes of molecular evolution

Greater fitness variation in asexual versus sexual populations suggests an underlying model in which asexual populations diverge genetically more rapidly and more stochastically than sexual populations. To directly evaluate this claim, we compared the number and spectrum of de novo mutations in sexuals and asexuals by sequencing adapted clones from generation 960 in each population. We first isolated two clones from each replicate population and assayed the fitness of each clone in both home and all away environments, finding that as expected these clone fitness measurements recapitulate the population-wide results described above (Fig. S3). We then conducted whole-genome sequencing of each clone at ∼20x depth, and used a standard bioinformatics pipeline to call mutations (Methods). Library preparation failed in three clones and we excluded six clones due to suspected whole-genome duplications in three populations that made mutation calling unreliable, leaving us with a total of 65 sequenced clones (for consistency, these three populations are excluded from both fitness and sequence analysis, see Methods).

We define a mutation as putatively fixed in a population if it is called in both clones isolated from that population, and as segregating if it is called in only one of these two clones. Using this definition, we find that asexual populations fix seven times as many mutations as sexuals, and contain twice as many segregating mutations (Fig. 2a; Generalized Wilcoxon test with Benjamini-Hochberg correction, both p < 0.001). We classified each mutation within a coding region as synonymous, nonsense, missense, or indel, and classified mutations in the latter three classes as ‘putatively functional’. We find that asexual populations contain three times as many putatively functional mutations as sexuals (Fig. 2a,b; Welch’s t-test, t = 10.20, p < 0.001). Thus, the average degree of genetic divergence of evolved clones from their ancestors is elevated in asexual compared to sexual lines, and this effect is even stronger among putatively functional mutations.

**Fig. 2.**
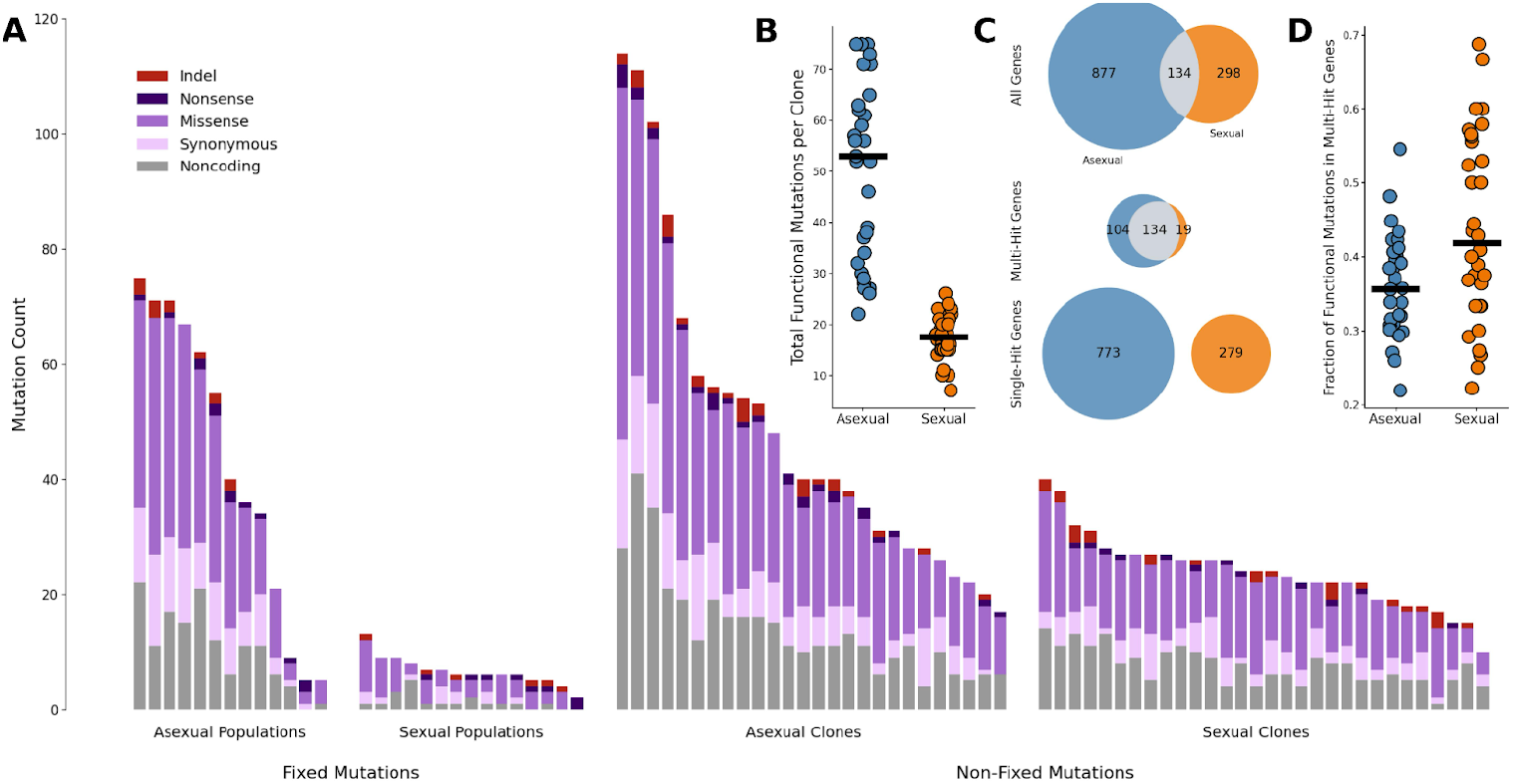
Recombination decreases the rate of genetic divergence and narrows the spectrum of mutational targets. **(A)** Number of fixed as well as non-fixed mutations is lower among sexual compared to asexual populations. **(B)** Distribution of total putatively functional mutations (non-synonymous SNPs + genic insertions and deletions) per clone, showing higher tendency and greater spread for asexual versus sexual clones. **(C)** Number of mutational targets (genes) with putatively functional mutations in sexual and asexual clones, broken down into multi-hit targets and single-hit targets. Genes are classified as multi-hit if they contain functional mutations in multiple replicate populations, and otherwise single-hit. **(D)** Fraction of putatively all functional mutations per clone found in multi-hit genes. Horizontal black bars indicate medians.

We classified a gene as ‘multi-hit’ if we identified a fixed or segregating functional mutation in that gene in multiple replicate populations, and as ‘single-hit’ otherwise. Because it is unlikely (though not impossible) for multiple independent hits to happen by chance, multi-hit genes are putative targets of adaptation. We find that both sexual and asexual lines contain mutations in a large number of these multi-hit genes (Fig. 2c), indicating that these putative targets of adaptation were similarly available to populations regardless of recombination regime. However, mutations in multi-hit genes represent a substantially larger proportion of all mutations in sexual compared to asexual clones (Fig. 2d; Welch’s t-test, t = -3.24, p = 0.002). In other words, molecular evolution in sexual populations involves accumulation of a higher proportion of putatively adaptive mutations than in asexuals.

### Hitchhiking genetic load leads to pleiotropic fitness costs in asexual lineages

Together, our fitness and mutation data support a model in which genetic linkage increases rates of genetic divergence in asexuals via the stochastic accumulation of non-target mutations. Such linked mutations constitute potential genetic load that can suppress fitness both in home and away environments and reduce the genotypic (and thus phenotypic) repeatability of local adaptation. To further explore this hypothesis, we sought to examine the relationship between fitness and the number of non-target mutations. Given that non-driver mutations (i.e., potential load) tend to outnumber beneficial driver mutations in mutational cohorts that segregate during adaptation (*14*), we predicted that clones with more overall putatively functional mutations should exhibit stronger direct and pleiotropic fitness liabilities.

Indeed, we found that asexual clones with relatively more putatively functional mutations had relatively lower fitness, and thus more genetic load, across both home and away environments (Fig. 3a). Using a linear model, we found that the relative fitness of asexually-evolved clones, in home and away environments, was negatively correlated with the number of mutations harbored by such clones (Fig. 3; linear regression, R^2^ = 0.52, F(1,27) = 29.19, p < 0.001). No such correlation was evident for sexually-evolved clones (Fig. 3; linear regression, R^2^ = 0.0005, F(1,26) = 0.01, p = 0.91). This pattern is strongest among single-hit mutations, which are likely enriched for non-targets, and weaker among multi-hit mutations, likely because they contain a smaller fraction of non-targets (Fig. S4; asexual single-hit R^2^ = 0.53, F(1,27) = 31.23, p < 0.001; asexual multi-hit R^2^ = 0.31, F(1,27) = 12.13, p = 0.002).

**Fig. 3.**
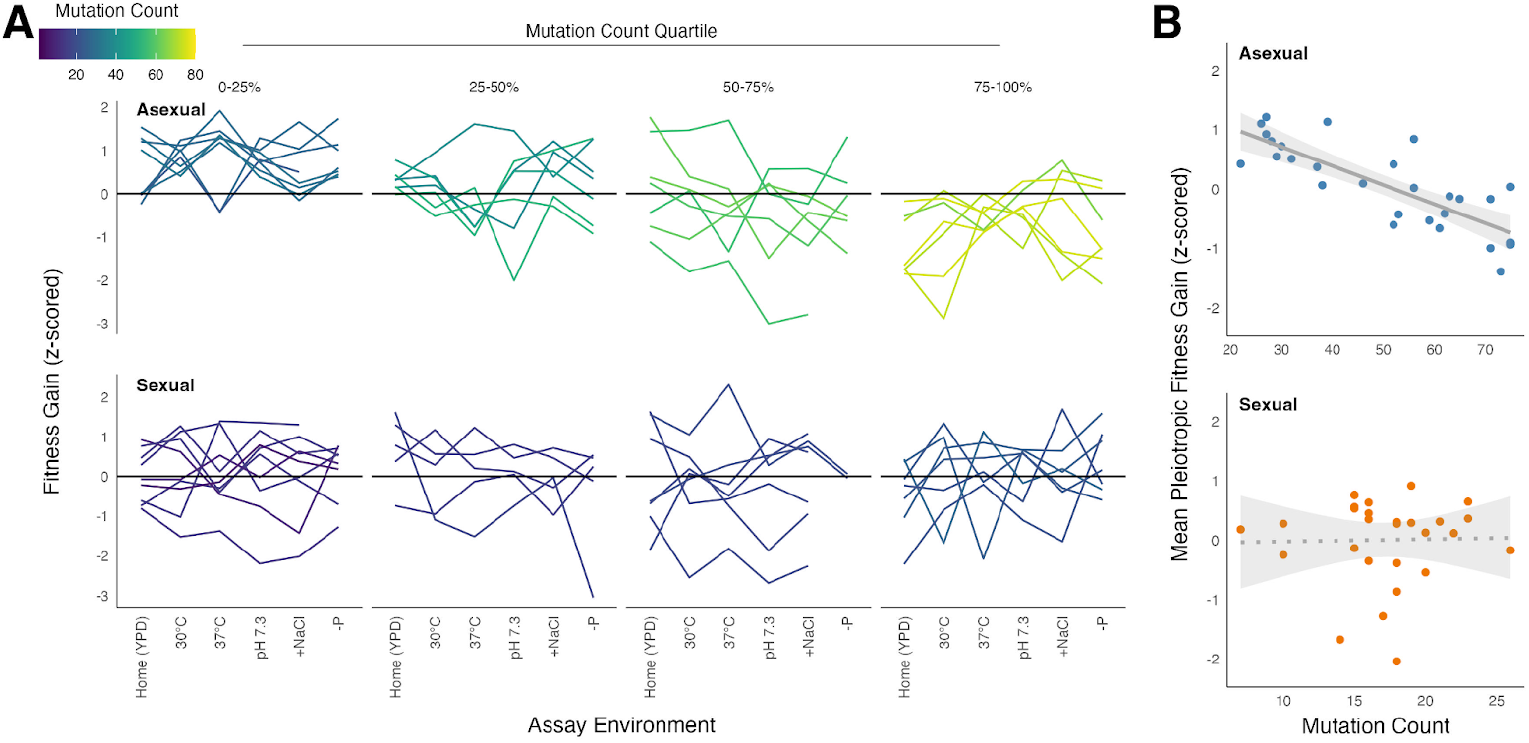
Asexual clones with more mutations show decreased pleiotropic fitness. **(A)** Fitness (scaled) across home and each away environments for clones binned by quartiles of total putatively functional mutations reveals asexual clones with more mutations are relatively less fit across all environments. **(B)** Asexual clones with more putatively functional mutations show lower relative mean fitness gain across away environments (i.e. pleiotropic fitness gain), whereas sexual clones show no such pattern.

### Backcrossing reveals that sex purges genetic load underlying pleiotropic costs of adaptation

To more directly test whether asexual lines harbor more load-causing mutations, we sought to separate load-causing from beneficial mutations by back-crossing adapted clones to an ancestral genotype. We predicted that parental genotypes with more genetic load would be more likely to produce back-crossed F1 offspring with higher fitness than their adapted parent, compared to parental genotypes with less genetic load. After isolating at least 60 F1 offspring from each of 19 back-crosses, we found that offspring of asexual parents were on average more fit than their parent in 6/10 crosses, while offspring of sexual parents were on average more fit than their parent in 2/9 cases (Fig. 4a,b). This difference is not statistically significant (Fisher’s exact test, p = 0.17), but the trend suggests that a single bout of sex (backcross) may be effective in purging genetic load underlying fitness costs for asexual lines in the home environment. In contrast, in sexual populations a single bout of sex with a non-adapted ancestor tends to be maladaptive, likely because these populations carry less load and hence the backcross purges more beneficial than deleterious mutations.

**Fig. 4.**
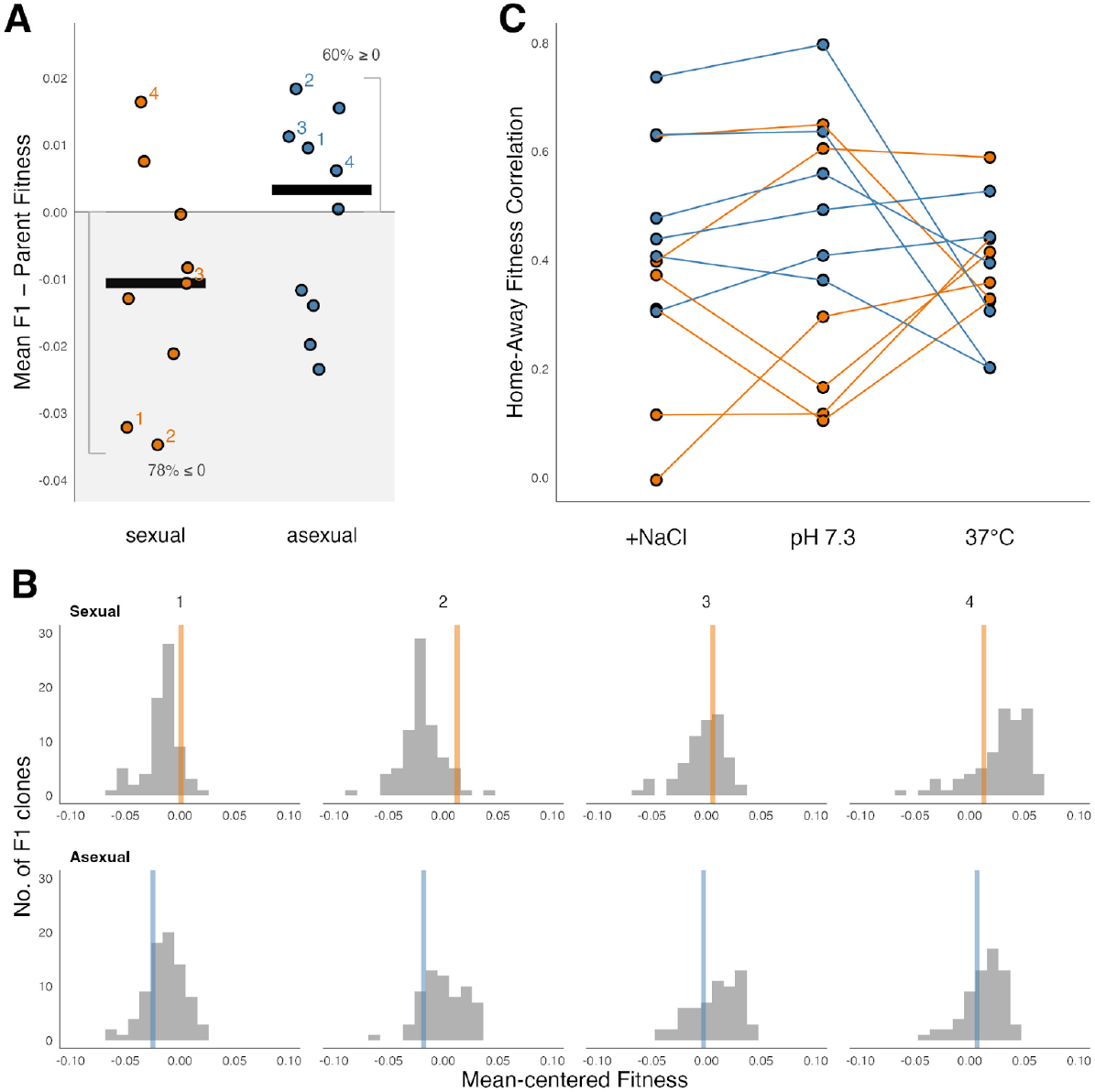
Ancestral back-crossing purges genetic load from adapted asexual parents in home and away environments. **(A)** Mean fitness difference in home environment (YPD) between parent and backcrossed offspring for sexually and asexually evolved parents (n ≥ 60 F1 clones assayed per cross). Horizontal lines indicate medians. Populations shown in (B) are indicated with numbers. **(B)** Examples of backcrossed F1 fitness distributions from 4 sexual (top) and 4 asexual (bottom) parents indicated in (A) with parental fitness of each strain indicated by a vertical bar. Fitness of parents and F1s are displayed mean-centered relative to mean of all parental strains (n = 19). Fitness values between -0.1 and +0.1 are shown for clarity, and these account for >99% datapoints. **(C)** Fitness correlation of backcrossed offspring across home and a subset of away environments (+NaCl, pH 7.3, 37°C).

We next used randomly selected subsets of our back-crossed F1 pools (6 asexual, 6 sexual) to more directly test the hypothesis that genetic load in the home environment underlies pleiotropic fitness costs in away environments. To do so, we assayed fitness of individual F1 offspring from back-crosses with adapted parents across a subset of our away environments (+0.2M NaCl, pH 7.3, and 37°C). Offspring fitness was positively correlated across environments (Fig. 4c): offspring with lower pleiotropic fitness also had lower fitness in the home environment, while relatively more fit offspring enjoyed this benefit across all away environments. Averaged across away environments, the relative fitness rank of an F1 genotype within its cohort was tightly positively correlated with its fitness rank in the home environment (Fig. S5; Spearman correlation ρ = 0.53-0.76 for sexuals and ρ = 0.62-0.91 for asexuals, all p <0.01 with Benjamini-Hochberg correction). This result reveals that genotypes with higher load in the home environment show similar extents of load in alternate environments, whereas more fit genotypes retain their advantage overall.

## Discussion

The evolution of fitness trade-offs across environments or ecological niches is central to local adaptation, ecological specialization, and eventually speciation (*15*). Reciprocal transplant experiments between natural populations of plants and animals from different environments have demonstrated that local adaptation is widespread in the wild, but that the magnitude of specialization is low (*16*). However, the factors underlying the speed, magnitude, and genetic basis of the evolution of specialization are not fully understood. Further, there are potentially key differences between the evolution of specialization in natural populations of sexual organisms and the asexual laboratory microbial populations often used to probe this question. Our results directly bridge this gap, by showing that specialization proceeds more rapidly and erratically in asexual lines, where genetic hitchhiking promotes specialization by increasing genetic load across environments. In contrast, in sexual populations, local adaptation leads to fewer pleiotropic fitness costs and hence enables lower-cost generalism to evolve.

This advantage of sex depends on the genetic basis of the pleiotropic costs of adaptation, which has traditionally been studied in the context of distinguishing antagonistic pleiotropy (*11*) from mutation accumulation (*17*–*21*). Earlier work has produced mixed evidence for both of these effects across many field and experimental studies (*22*–*40*). Our results are consistent with mutation accumulation being dominant over antagonistic pleiotropy, since recombination purges accumulated mutational load and alleviates pleiotropic fitness costs. If antagonistic pleiotropy were strong, we would expect sexual lines to have lower fitness than the ancestor in away environments, which they do not. Moreover, we see no antagonistic pleiotropy even among asexuals after a single bout of sex, with fitness in the home and away environments positively correlated. This suggests that, at least in our system, direct biochemical or physiological tradeoffs are significantly less important than hitchhiking deleterious load in driving specialization.

A longstanding problem in evolutionary biology is the maintenance of sex in almost all eukaryotes despite its clear costs to survival, including the costs of mating, increased risks of predation or parasitism, and breaking up coadapted gene complexes or ‘winning genotypes’ (*1*– *4, 6*). Our work lends further support to earlier studies demonstrating that recombination speeds adaptation by rescuing beneficial driving mutations from deleterious backgrounds (‘ruby in the rubbish’) (*5*). This is one important factor explaining why sex is so pervasive. Additionally, we show here that this genetic load is responsible for pleiotropic fitness tradeoffs in asexual populations, which are largely alleviated by recombination. In other words, sex not only speeds local adaptation but it also reduces its pleiotropic costs. This effect of reducing the costs of generalism or niche expansion may help explain why sex is prevalent over longer evolutionary histories, where environmental conditions change and fluctuate.

Much earlier work has argued that the evolution of evolvability, often defined as the ability to generate adaptive variation, is an important response to changing environmental conditions (*41*– *43*). However, empirical demonstrations of this effect have primarily been limited to asexual microbes (*44*–*46*). In contrast, in sexual populations, recombination is predicted to constrain the evolution of evolvability by separating loci that increase evolvability from the adaptive variation they generate (*47, 48*). Our results here offer a simpler explanation for how sexual populations can adapt in the face of new or changing environments. Indeed, we find that recombination increases the robustness of the evolutionary response to shifting conditions, despite (and indeed because of) the fact that these populations generate less genetic variation than asexuals.

Further work is needed to fully understand the interplay between sex, pleiotropy, and ecological generalism. A more direct approach to demonstrate that pleiotropic fitness costs are caused by costly genetic load would be to reconstruct individual putative beneficial drivers and deleterious hitchhikers to directly measure their fitness effects. While technically challenging to conduct at scale, this is now becoming feasible using CRISPR-based methods. It would also be interesting to analyze adaptation on standing variation and in diploids rather than haploids, and to test whether these benefits of sex are diluted or amplified in such scenarios. Finally, much work on antagonistic pleiotropy has focused on life history trait tradeoffs, such as between pre- and post-reproductive survival, conflict between the sexes, and even between asexual and sexual life cycle phases (*49, 50*). Future work considering this richer set of important phenotypes, expanding our approach to other facultatively sexual species with varying life histories, and integrating these studies with high-throughput phenotyping, genotyping, and genetic manipulation techniques may enable us to shed even more light on the critical question of why sex is so widespread across biology.

## Materials and Methods

### Experimental design

#### Strains

We established independent sexual and asexual lines (n = 17 and 18 respectively) from single colonies of ancestral haploid *MAT****a*** and *MATα* W303 parent strains, MJM64 and MJM36 respectively (*5*). These strains are isogenic apart from several selectable markers that allow us to control mating and recombination. Specifically MJM64 has genotype *MAT****a****-KanMX, ho, Pr*_*STE5*_ *-URA3, ade2-1, his3*Δ::*3xHA, leu2*Δ::*3xHA, trp1-1, can1::Pr*_*STE2*_ *-HIS3-Pr*_*STE3*_ *-LEU2* and MJM36 has genotype *MATα-HphMX, ho, Pr*_*STE5*_ *-URA3, ade2-1, his3*Δ::*3xHA, leu2*Δ::*3xHA, trp1-1, can1::Pr*_*STE2*_ *-HIS3-Pr*_*STE3*_ *-LEU2*. The key feature of these strains that permits controlled mating and recombination is that each mating type harbors a drug marker tightly linked to the mating type locus (KanMX for *MAT****a***; HphMX for *MAT****α***), along with nutrient markers under the control of mating-type specific promoters (Pr_*STE2*_ -HIS3 for *MAT****a***; Pr_*STE3*_ -LEU2 for *MATα*) and the counterselectable URA3 marker under the control of a haploid-specific promoter (Pr_*STE5*_ -URA3). These markers permit selection using at least two markers for *MAT****a*** haploids, *MATα* haploids, and diploids, which we have shown in previous work is essential for maintaining efficient sexual cycles through long-term laboratory evolution (*5, 12*).

#### Propagation regime

All yeast lines in this study were propagated using our previously described protocol (*12*); all methods for serial propagation, archiving, contamination checks, and steps of the sexual cycle were conducted exactly as previously described. Key features of the experimental protocol are illustrated in Fig. S1. Briefly, each population consists of two paired lines, which we passaged using a standard asexual batch culture protocol in 128μL of rich laboratory media (YPD; 1% yeast extract, 2% peptone, 2% dextrose) in unshaken flat-bottom 96-well microplates, with a daily 1:2^10^ dilution conducted using a Biomek FX liquid handling robot (Beckman Coulter). Each sexual population consisted of one *MAT****a*** and one *MATα* line, while each asexual population consisted of two *MAT****a*** lines. Every 120 generations, we implemented sexual cycles in sexual populations, and for consistency “mock” sexual cycles in asexual populations. Using this approach, we evolved each population for a total of 960 generations, with a total of 8 bouts of recombination (or mock treatment, for the asexual lines) throughout the experiment.

In each sexual cycle, we mixed saturated culture from each *MAT****a*** and *MATα* population pair in fresh YPD with a 1:10 dilution and allowed these cells to mate for 6 h at 30°C in unshaken 96-well plates. We then diluted the mating culture 1:10 into 128 μL of diploid selection media (YPD + hygromycin [300 μg/mL] + G418 [200 μg/mL]), which only permits growth of diploids that contain both of the separate drug markers linked to each mating type locus. After overnight growth in diploid selection media, we diluted cultures 1:10 into 128 μL sporulation media (Spo++, 0.25% yeast extract, 1.5% potassium acetate, 0.05% glucose, 1X CSM amino acid stock) and held cultures with shaking (1,050 rpm) in 96-well plates for 2 d at 30°C. Following sporulation, we disrupted tetrads by incubating cultures in a suspension of glass beads containing zymolyase (20 μg/mL) and shaking the plates for 1 hour at 37°C. To isolate spores into haploids of distinct mating types, we diluted the sporulation culture 1:10 into *MAT****a*** selection media (CSM −uracil −histidine +G418) or *MATα* selection media (CSM −uracil −leucine +hygromycin) and allowed cultures to saturate overnight at 30°C. This re-establishes the pair of lines, one *MAT****a*** and the other *MATα*, after which we can resume normal batch culture propagation.

For consistency, our “mock” sexual cycles for asexual lines were designed to be as similar as possible to the protocol above, without actually implementing recombination. Specifically, for each asexual population we combined the two paired *MAT****a*** lines and selected in the appropriate single-drug media. Each line was then held in sporulation media for 2 d following identical dilution procedures as for the sexual lines, and then propagated in *MAT****a*** selection media as described above for the sexual cycle. This ensures that the asexual lines experience the same bottlenecks and effective population sizes as sexual lines, and as close to identical selection pressures during the sexual or mock sexual cycles as possible.

One important caveat is that because the asexual populations consist of two paired *MAT****a*** lines, while the sexuals consist of one *MAT****a*** and one *MATα* line, there could be differences arising from differences in the evolution of these two mating types (e.g. arising from somewhat different mutation rates between mating types). Some such differences have been observed in related strains in earlier work, with *MAT****a*** strains accumulating mutations at a somewhat higher rate than *MATα* lines (*51*). However, since both sexual and asexual populations include evolution of at least one *MAT****a*** line, and all fitness assays and sequencing are conducted in these *MAT****a*** lines, this effect can at most represent a modest difference in effective population size, which cannot explain the dramatic differences we observe between evolution in sexual versus asexual populations.

### Fitness assays

At generation 960, following the 8th sexual (or mock-sexual) cycle, we measured the fitness gain of whole populations relative to the ancestor of all evolved populations. Fitness assays were all conducted on the *MAT****a*** line from each sexual population, or one of the two *MAT****a*** lines (chosen at random) for each asexual population. Procedures for fitness assays closely followed published protocols (*12, 52*) and entails serial propagation in YPD as described above for 2 dilution cycles (approximately 20 generations). By seeding a diploid fluorescent reference strain (yKK433) into the starting populations, we track the change in frequency of the fluorescent reference subpopulation compared to the test population by scoring ∼10-20,000 cells via flow cytometry. This reference was constructed by mating DivAncCit (*53*) and yAN433 (*51*), producing genotype *MAT****a****/MATα his3*Δ::*HIS3-ymCitrineM2331/his3*Δ, *LYS2/lys2*Δ, *leu2-3,112/leu2*Δ, *ade2-1/ADE2, trp1-1/TRP1, ura3::Pr*_*FUS1*_ *-yEVenus/ura3*Δ, *bar1*Δ::*ADE2/BAR1, hmlα*Δ::*LEU2/HMLα, gpa1*Δ::*GPA1-G1046T-NatMX/GPA1, RME1/Pr*_*RME1*_ ::*ins-308A, YCR043C/YCR043C*Δ::*HphMX4, CAN1/can1*Δ::*Pr*_*RPL39*_ *-ymCherry-Pr*_*STE2*_ *-SpHIS5-Pr*_*STE3*_ *-LEU2*. Because this reference strain is more fit than most evolved lines, we typically seeded it at an initial frequency of 10-20%, to ensure that test populations with relative fitness defects of up to 20% compared to the reference could be accurately measured. Prior to all fitness assays, test and reference populations were thawed on the benchtop, aliquoted (4 μL) into 128 μL fresh YPD, and allowed to saturate overnight at 30°C. For all assays, we randomized between three and four independent replicate assays per population per assay environment to positions within and between 96-well fitness assay plates, to avoid confounding with spatial and/or plate-level effects. We estimated fitness by calculating the per-generation increase in relative frequency of the test strain between final and initial time points, on the logit scale, as ŝ = Δt^−1^ [logit(*f*_*2*_) − logit(*f*_*1*_)], where logit(*f*_*t*_) = ln[*f*_*t*_ /(1 − *f*_*t*_)]. Estimates of *f*_*t*_ for test populations were taken as the fraction of non-fluorescent events per time point after manually gating fluorescent cells using FlowJO. All wells with fewer than 1250 events of either the test or reference population by *t*_*2*_ were excluded from analyses, and |ŝ| estimates more extreme than 20% were considered outliers and were removed from all analyses on the basis of being biologically implausible and difficult to measure accurately.

To make estimates of fitness at the individual genotype level, we isolated adapted clones from each of the sexual and asexual populations by dilution plating population aliquots onto *MAT****a*** selective media. After 2 d of growth at 30°C, we randomly isolated four clones per line by picking colonies whose center was most adjacent to the corners of a superimposed grid. Clones were grown overnight to saturation in YPD, re-arrayed in a single 96-well plate, and frozen in 25% glycerol.

For measurements of fitness in the five ‘away’ environments, we conducted fitness assays exactly as described above, but in separate media. We used the same synthetic complete base medium (0.2% SC, Sunrise Science, 0.671% yeast nitrogen base + ammonium sulfate [YNB+N], 2% dextrose) for all away environments but varied the temperature at which cultures were incubated (30°C or 37°C), or the presence of osmotic stress (+0.2 M NaCl), a buffering agent (to pH 7.3 with PIPES buffer, piperazine-*N,N*′bis[2-ethanesulfonic acid]), or phosphate limitation (with yeast nitrogen base + ammonium sulfate - potassium phosphate in place of YNB+N, MP Biomedicals). Fresh media was prepared from concentrated stocks held at 4°C prior to each fitness assay.

### Whole-genome sequencing and Variant Calling

We sequenced our set of n = 74 adapted clones by generating libraries from DNA extracted with a PureLink Pro 96 Genomic DNA Purification Kit, following previous work (*54*). We conducted 2×150 bp paired end sequencing on the Illumina HiSeq 5000 platform and with expectation of 10–20X coverage for all genomes. Libraries with fewer than 10X average read coverage were resequenced, and sequencing data was pooled prior to mutation calling. All libraries were prepared with dual-indexing using standard Nextera primers (*54*).

For sequencing analysis, we first created a new ancestral reference genome to correct for any mutations present in the ancestral strains compared to a previous *S. cerevisiae* W303 genome (*55*): by pooling read alignments from all 64 clones, any variants called between this pool and the reference are assumed to have been present in our ancestor. Specifically, for each clone, we trimmed Illumina reads with NGMerge (*56*), aligned them to the W303 reference genome using BWA (*57*), and marked duplicate reads with Picard (*58*). We then used Samtools (*59*) to combine these alignments, and Pilon (*60*) to polish the W303 reference with any variants called from this pool, producing our corrected ancestral reference genome. We then repeated the trimming, alignment, and duplicate marking steps for all clones using this new corrected reference, and called variants using GATK (*61*) (in particular, HaplotypeCaller, GenomicsDBImport, and GenotypeGVCFs). We annotated these called variants with SnpEff (*62*).

We filtered the list of variants by excluding all occurring in telomeres, since alignment errors and repetitive sequences make variant calling difficult in these regions, and in mitochondria, since these are inherited uniparentally. We discarded variants with low support: either if the locus had overall <10x average coverage across clones (putatively areas where sequencing or alignment is challenging), or if the specific variant had <5 alternate allele reads. Further, we removed variants that appeared in more than one population, under the assumption that exact nucleotide-level parallelism across populations is negligibly rare in our experiment [only one example was seen in 10,000 generations of asexual experimental evolution previously (*51*)], and these are instead more likely to be alignment errors [confirmed by manual inspection of BAM files using IGV (*63*)]. Finally, we excluded variants with an alternate allele frequency <0.75, which again were more often alignment errors. Fixed mutations were estimated as the intersection of the mutation sets from the two sequenced clones isolated from each adapted population.

Library preparation failed in three asexual clones. For six further clones from three populations (two sexual and one asexual), we identified potential whole-genome duplication events from our sequencing data, based on finding alternate allele frequency distributions centered around 50% (Fig. S6). Since this makes mutation calling unreliable, we also excluded these from all sequence and fitness analyses for consistency, leaving us with a total of 65 sequenced clones from 32 populations.

### Ancestral back-crossing

#### Crossing regime and F1 isolation

We randomly selected 9 sexual and 10 asexual parental lines and backcrossed one of the clones isolated from each line (see above) to the *MATα* ancestor. Specifically, we performed mating in a total volume of 300 μL in deep 96-well plates with the same steps as the sexual cycle. For sporulation, we waited up to 4 d in sporulation media to maximize the sporulation fraction. Following asci disruption via zymolyase treatment, we resuspended spores in *MAT****a***-selective media and allowed them to recover for 3 h prior to plating on *MAT****a*** selection plates. Following this, we replica-plated once again onto the same media type and allowed cultures to grow for up to 2 d at 30°C. We then systematically isolated at least 64 F1 clones per backcross by picking colonies whose centers were proximal to nodes of a superimposed lattice. In some cases, F1 densities were too high to confidently pick single colonies; in these cases, plates were scraped, re-suspended, and re-plated at higher dilution onto *MAT****a*** selection plates.

#### F1 fitness assays

We measured fitness of these backcrossed F1 offspring in the home environment YPD using the protocol described above. For two of the asexual parental clones, only n=16 and n=39 of the F1 fitness assays were successful respectively, but all others had at least n=60 F1s successfully assayed. In parallel, we assayed 6 asexual and 6 sexual F1 offspring sets (n=24-80 F1s) across three of our five away environments (37°C, +0.2 M NaCl, pH 7.3). F1 clones and their parents (i.e. evolved sexual or asexual lines) were assayed in the same plates and spread across several batches, with a single assay replicate per F1 genotype and 2-4 parental replicates per plate per environment.

## Acknowledgments

We thank the Bauer Core facility at Harvard and members of the Desai laboratory for experimental assistance and comments on the manuscript. Computational work was performed on the FASRC Cannon cluster supported by the FAS Division of Science Research Computing Group at Harvard University.

## Author contributions

Conceptualization: SVP, PTH, MMD

Formal analysis: SVP, PTH

Funding acquisition: MMD

Investigation: SVP, PTH, CS, KK, AL

Methodology: SVP, PTH, CS, KK, AL

Resources: MMD

Supervision: MMD

Visualization: SVP, PTH

Writing – original draft: SVP, PTH, MMD

Writing – review and editing: SVP, PTH, CS, KK, AL, MMD

