## Supplementary Figures for "Sex decreases the pleiotropic costs of local adaptation"

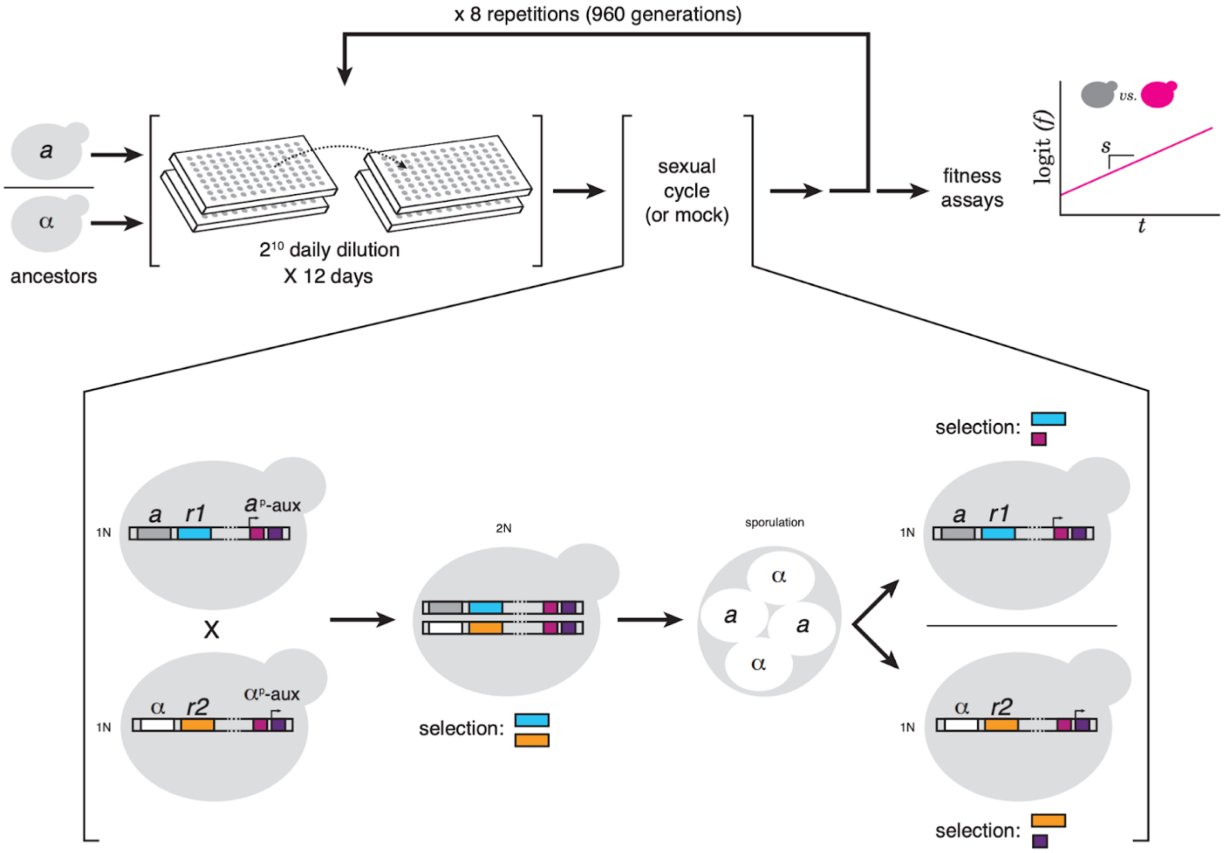


***Fig. S1. Schematic of experimental procedures and genetic system permitting controlled mating and recombination of haploid yeast lines***

*Sexual (n=17) and asexual (n=18) populations are propagated using standard asexual batch culture in 96-well plates for 960 generations with or without sexual cycles every 120 generations, respectively. Mating type loci are linked to drug resistance markers (r1, blue, & r2, orange) and auxotrophic markers (pink, purple) under control of mating type-specific promoters, enabling double drug-auxotrophic selection post sporulation. After 960 generations of propagation, populations are assayed for fitness using flow cytometry relative to a fluorescent reference strain.*

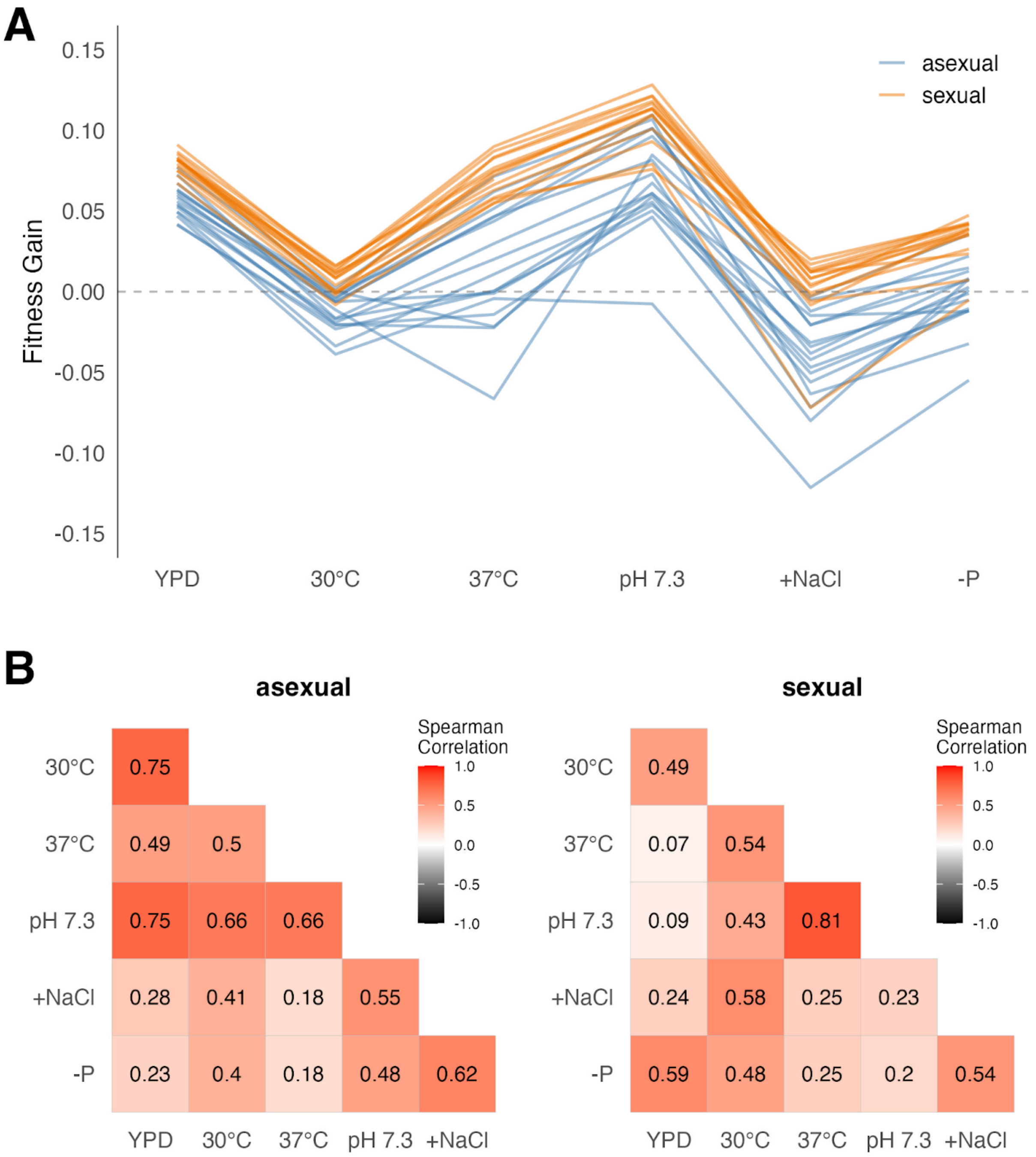


***Fig. S2. Antagonistic pleiotropic tradeoffs are absent in both sexually and asexually evolved populations***
***(A)*** *Distribution of fitness changes relative to the ancestor across home (YPD) and away environments (SC 30°C, 37°C, pH 7.3, +NaCl, -phosphate), with each independent sexual or asexual population connected across environments by horizontal lines.* ***(B)*** *Pairwise Spearman correlation in fitness gain between all environments for asexual and sexual populations.*

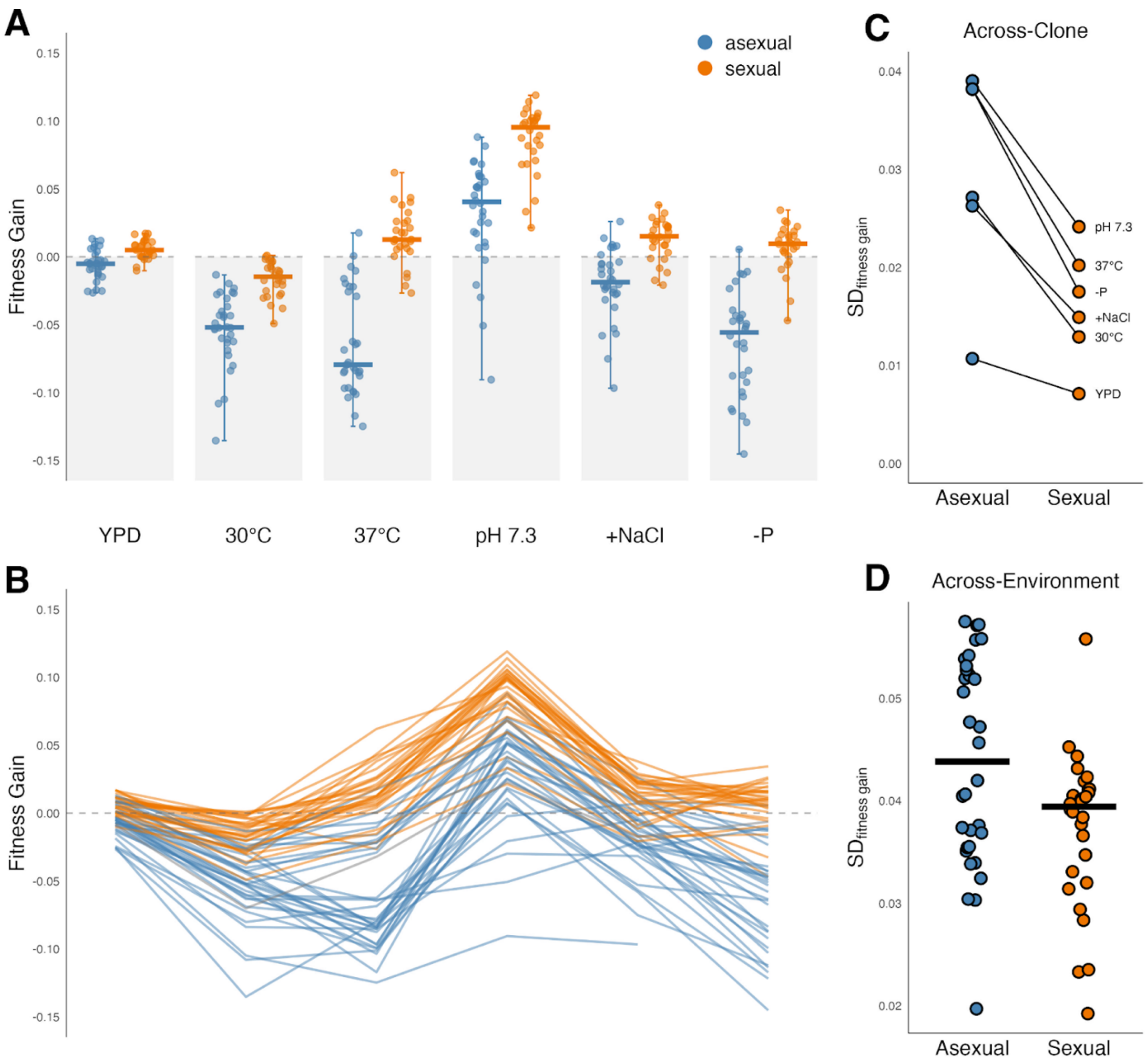


***Fig. S3. Recombination increases fitness gain, repeatability, and decreases pleiotropic costs of adaptation in clones***

***(A)*** *Distribution of fitness changes relative to the ancestor across home (YPD) and away environments (SC 30°C, 37°C, pH 7.3, +NaCl, -phosphate), with each independent sexual or asexual clone represented.* ***(B)*** *Distribution of fitness changes across environments with each clone represented by a horizontal line.* ***(C)*** *Standard deviation in fitness across replicate clones in asexual compared to sexual populations in home and each away environment.* ***(D)*** *Standard deviation in fitness of individual clones across environments (crossbars indicate medians).*

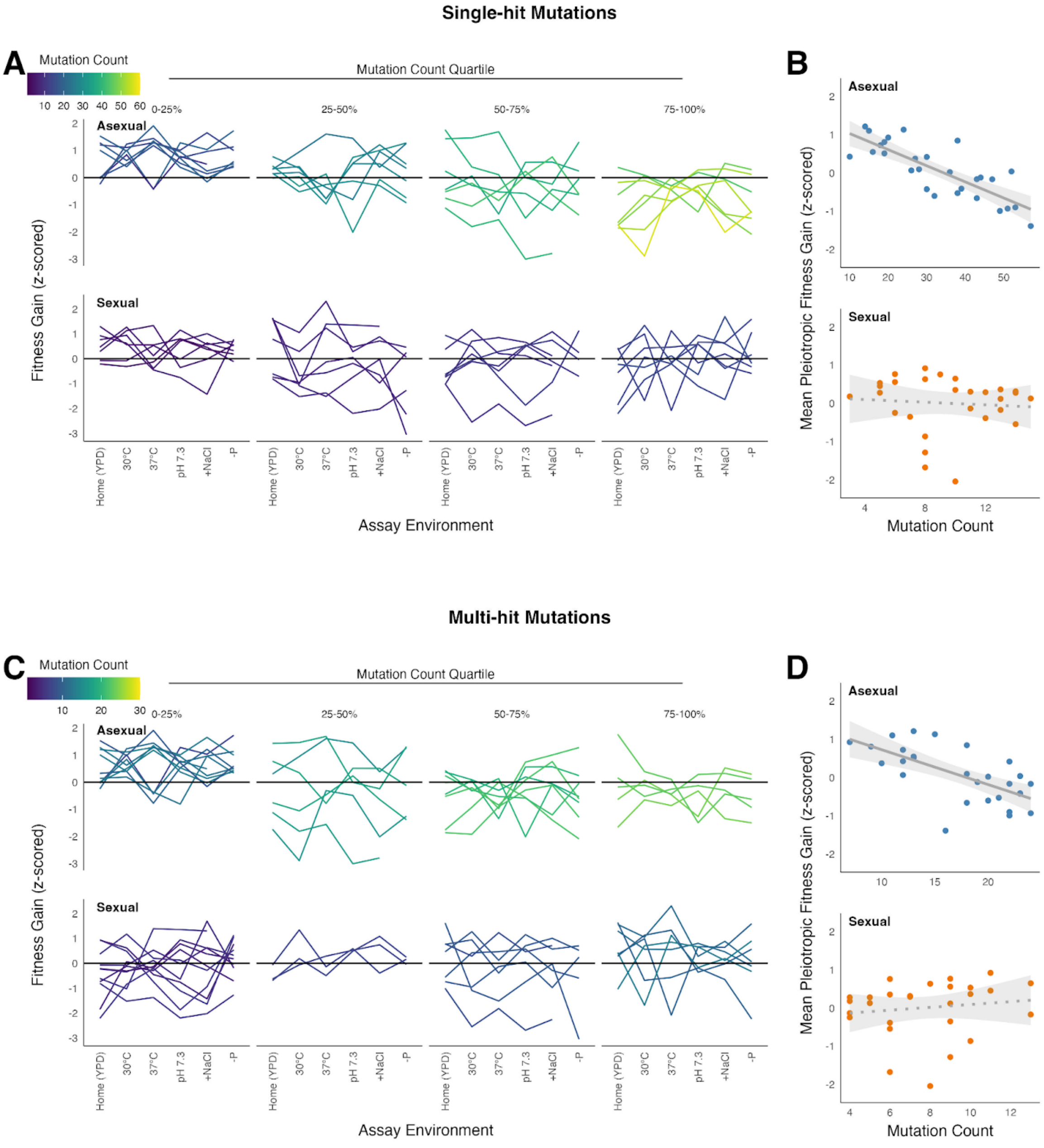


***Fig. S4. Asexual clones with more single-hit and multi-hit mutations show decreased pleiotropic fitness***

*Identical to Fig. 3, but subsetted to only functional mutations in single-hit genes (A, B) or multi-hit genes (C, D)*

***(A, C)*** *Fitness (scaled) across home and each away environments for clones binned by quartiles of putatively functional mutations reveals asexual clones with more mutations are relatively less fit across all environments.* ***(B, D)*** *Asexual clones with more putatively functional mutations show lower relative mean fitness across away environments, whereas sexual clones show no such pattern.*

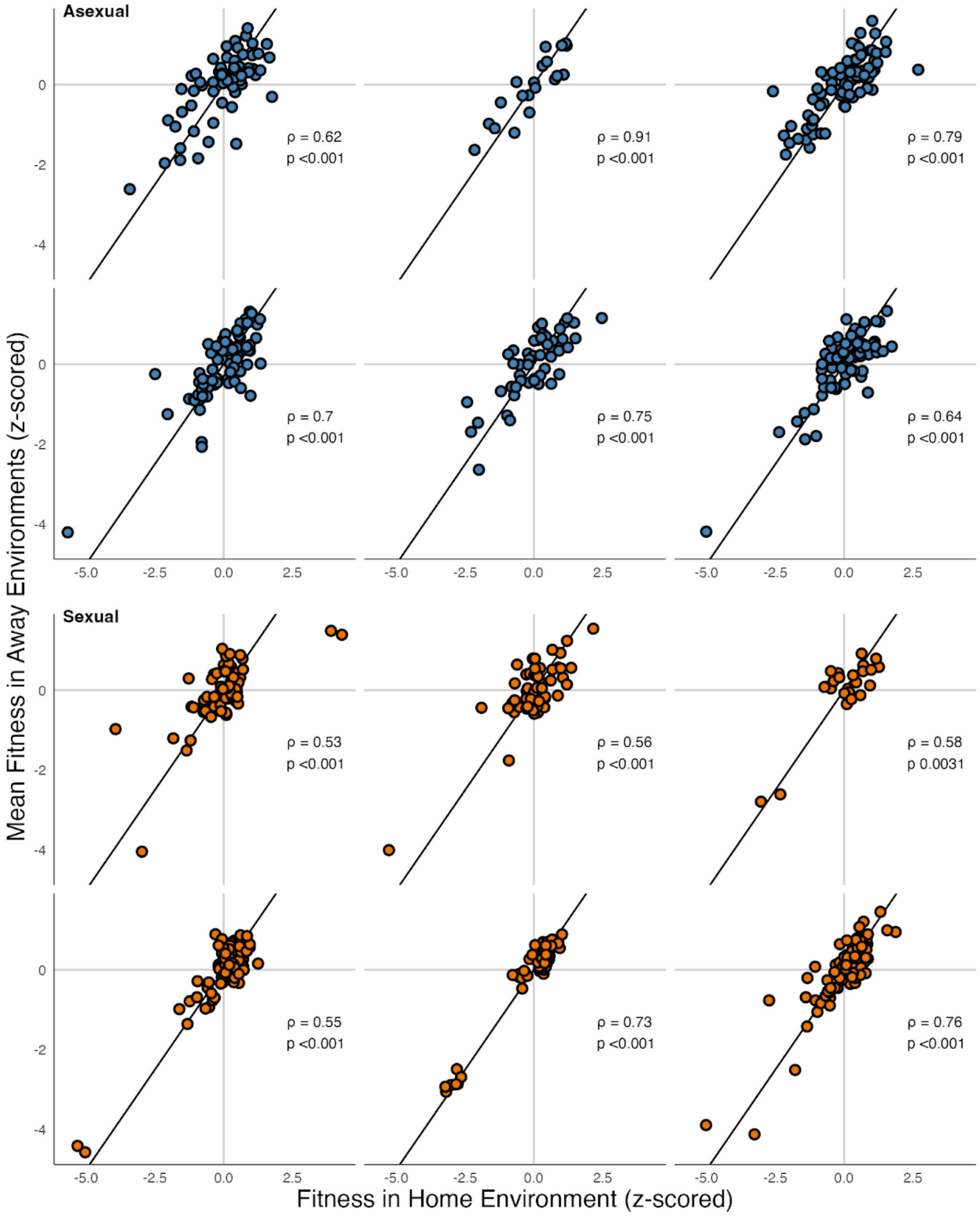


***Fig. S5. Home and pleiotropic fitness are positively correlated for asexual and sexual backcrossed offspring***

*Scatterplots showing positive correlations between home (x) and mean z-scored away fitness (y) for pools of backcrossed offspring. Each plot corresponds to one parental evolved clone. Diagonal indicates 1:1 line. P values adjusted for multiple testing with Benjamini-Hochberg correction.*

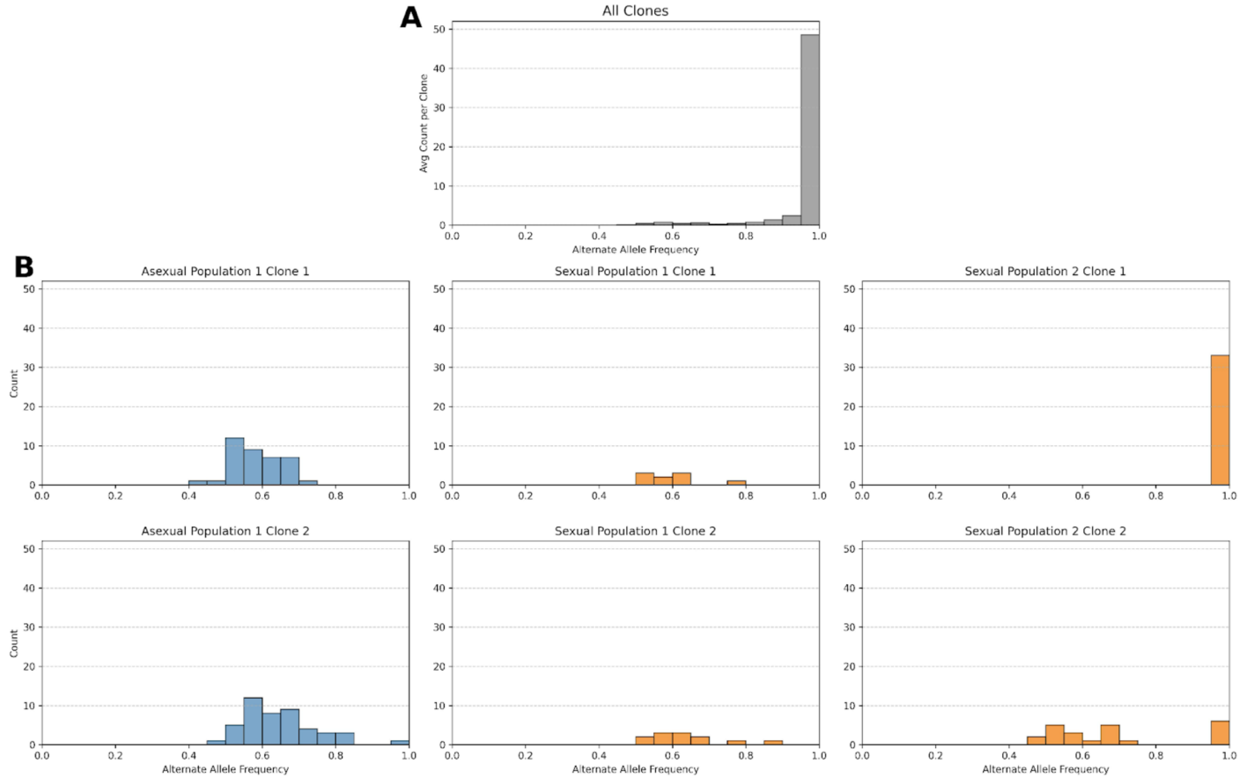


***Fig. S6. Alternate allele frequency distributions for clones with suspected whole-genome duplications***

***(A)*** *Histogram of alternate allele frequency per variant averaged across all clones.* ***(B)*** *Histograms of alternate allele frequency per variant for 6 clones (2 asexual, 4 sexual) from populations with suspected whole-genome duplications, evidenced by alternate allele frequency distributions centered around 0.5 (compared with overall distribution centered close to frequency 1).*
